# Human chaperones untangle fibrils of the Alzheimer protein Tau

**DOI:** 10.1101/426650

**Authors:** Luca Ferrari, Willie J.C. Geerts, Marloes van Wezel, Renate Kos, Aikaterini Konstantoulea, Laura S. van Bezouwen, Friedrich G. Förster, Stefan G.D. Rüdiger

**Affiliations:** Cellular Protein Chemistry, Bijvoet Center for Biomolecular Research, Utrecht University, Padualaan 8, 3584 CH Utrecht, The Netherlands; Science for Life, Utrecht University, Padualaan 8, 3584 CH Utrecht, The Netherlands; Cryo Electron Microscopy, Bijvoet Center for Biomolecular Research, Utrecht University, Padualaan 8, 3584 CH Utrecht, The Netherlands; Utrecht University of Applied Sciences, Heidelberglaan 7, 3584CS Utrecht, The Netherlands

**Keywords:** Proteostasis, Protein folding, Protein quality control, Neurodegeneration, Disaggregase, Hsp70

## Abstract

Alzheimer’s Disease is the most common neurodegenerative disorder. A hallmark of this disease is aggregation of the protein Tau into fibrillar tangles, which is ultimately linked to neuronal death ^1,2^. Oligomeric precursors of Tau fibrils are suspected to be the neurotoxic agent while fibrils themselves may be less harmful end products of the aggregation process ^3,4^. Evolutionary conserved families of molecular chaperones maintain protein homeostasis in healthy cells, preventing aggregation ^5,6^. Here, we investigate whether such chaperones could possibly reverse the aggregation reaction and dissolve Tau fibrils. Indeed we find that the human Hsp70 chaperone system disaggregates Tau fibrils. Both the bacterial and human Hsp70 chaperone systems disassemble fibril superstructures assembled of several fibril strands into single fibrils, indicating that this is an evolutionary conserved capacity of the Hsp70 system. However, further disaggregation of Tau fibrils into oligomers and even monomers is reserved to the human homologue. Thus, although bacteria possess an effective machinery to dissolve amorphous aggregates ^7-9^, we see that they do not have the means to disaggregate fibrils. Fibrillar aggregates, therefore, require different chaperone systems than amorphous aggregates, and this is a property acquired by Hsp70 during evolution. This makes the Hsp70 system an interesting target for novel drug strategies in Alzheimer.

## INTRODUCTION

Tau is an intrinsically disordered protein of 441 residues in its longest isoform ^3,10^. Its function is the stabilisation of microtubules, mediated by its repeat domain (Tau-Q244-E372; Tau-RD ^11^). Tau interaction with microtubules is controlled by phosphorylation, and in healthy neurons Tau is continuously degraded ^12^. Formation of Tau fibrils is regarded as an irreversible process linked to the neuronal death in Alzheimer’s disease, in which oligomeric intermediates are suspected as actual toxic species ^2-4^ The neurotoxic effect of Tau aggregation is not depending on a specific form of aggregates. Tau fibrils in Alzheimer form dimeric straight filaments and paired filaments ^13^. Tau fibrils in Pick’s disease contain only three of the four repeats of the repeat domain and form monomeric fibrils with entirely different side chain pairing than the Alzheimer fibrils ^14^. Tau-RD with the pro-aggregating mutation ΔK280 but lacking the five residues of the Alzheimer fibril aggregates even faster than the full-length protein and constitutes an established disease model, causing neuronal death and memory loss in mice ^15^.

It is crucial for longevity of neurons to prevent any form of Tau aggregation. The molecular chaperones Hsp70 and Hsp90 are involved in controlling degradation of Tau, preventing uncontrolled appearance of free Tau ^16-19^. This process is disturbed when Tau fibrils form in Alzheimer’s Disease. The role of chaperone control in the later steps of Tau aggregation, however, is poorly understood.

We wondered whether molecular chaperones could be capable to reverse aggregation of Tau. Recently it became clear that metazoan Hsp70 chaperone system has disaggregase activity ^20^. Hsp70 chaperones are ATP-dependent machines that are regulated by ATPase-stimulating J-proteins and nucleotide exchange factors. A trio consisting of the Hsp70 homologue Hsc70, the J-protein Hdj1 and the nucleotide exchange factor Apg2 disaggregates α-synuclein fibrils to monomers ^21^. Hsc70 is the housekeeping Hsp70 chaperone and under normal conditions the most important cytosolic Hsp70 protein. Therefore, we set out to test whether the human Hsc70 system can disaggregate Tau fibrils.

## RESULTS

For investigating disaggregation of Tau fibrils, we used the pro-aggregating ΔK280 variant of Tau-RD, with a FLAG tag to facilitate detection (Tau-RD*), which fibrillates more aggressively than the wildtype. It contains both Hsp70 binding sites and the largest part of the Tau fibril structure ^13, 22,23^. The lack of the highly charged and less hydrophobic N-terminus facilitates aggregation. We used the established heparin protocol to initiate fibril formation ^24^. We confirmed by transmission electron microscopy (TEM) that Tau-RD* indeed forms fibrils with regular periodicity of 50-100 nm (**Fig. 1a** and **Fig. S1b**). Thus, we produced fibrils of this neurotoxic fragment to analyse disaggregation by chaperones.

**Figure 1.**
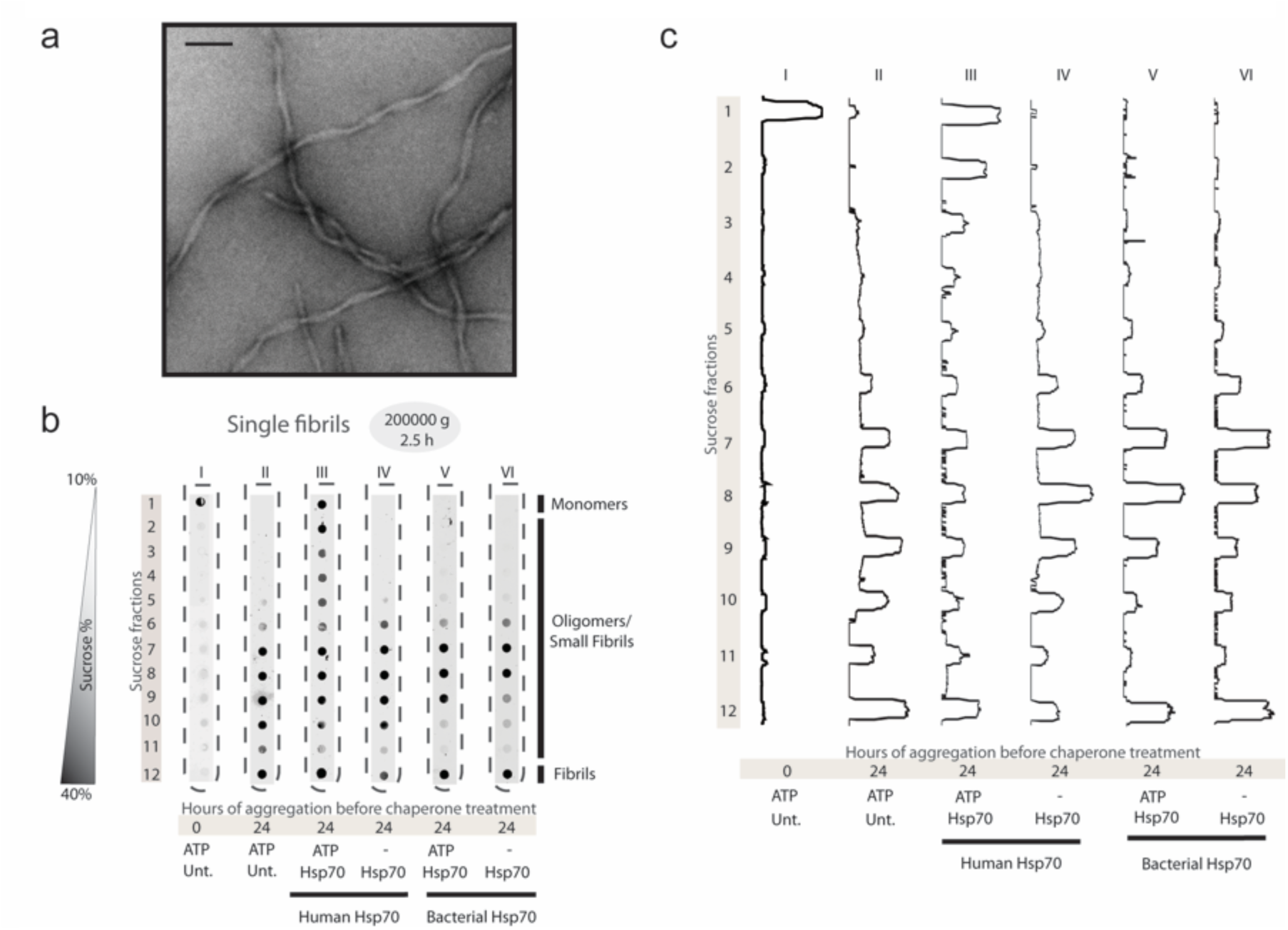
Only human Hsp70 disaggregates Tau single fibrils. **a**. TEM of Tau single fibrils. Scale bar 100 nm. **b**. Density gradients of preformed, heparin-induced Tau-RD* single fibrils, untreated (Unt.) or treated for 24 h with chaperones as indicated. Centrifugation was performed at 200,000 g for 2.5 h. **c**. Intensity profiling of (**b**), normalised to a 0 to 1 scale.

To monitor aggregation, we separated monomers, oligomers and fibrils mixtures of Tau-RD* on density gradients followed by dot blotting detection with a fluorescently labelled antibody. Non-aggregated Tau-RD* sediments as monomers on top of the gradient (**Fig. 1b**, tube **I**, fraction 1). After induction of aggregation by heparin, the sample does not contain monomers anymore. Instead we observe a mixture of soluble oligomers in the middle of the gradient (**Fig. 1b**, tube **II**, fractions 6-11) while fibrils move to the bottom (tube **I**, fraction 12).

We now tested the potential of the human Hsc70 machinery to disaggregate Tau fibrils. We used heparin-aggregated Tau-RD* and added the chaperone system after fibril formation. Strikingly, the density gradient analysis revealed that Hsc70 generated monomers (**Fig 1b**, tube III). The Hsc70 disaggregation activity was strictly ATP-dependent (**Fig 1b**, tubes III and IV), indicating that disassembly of Tau fibrils works along the same mechanistic lines as other activities of Hsc70 in folding, unfolding and disassembly. The intensity profile of the dot blots visualised relative changes in the size distribution of Tau-RD* (**Fig. 1c**). In the presence of the Hsc70 and ATP, the most prominent appearance of Tau-RD* is in fractions 1 and 2 (**Fig. 1c**, profile III), corresponding to monomers and small oligomers. After Hsc70 action we did not observe defined particles anymore by TEM and residual protein material became positively stained (**Fig. S1a**). Thus, fibrils and larger oligomers are not inert in the presence of a functioning Hsc70 system.

**Figure 2.**
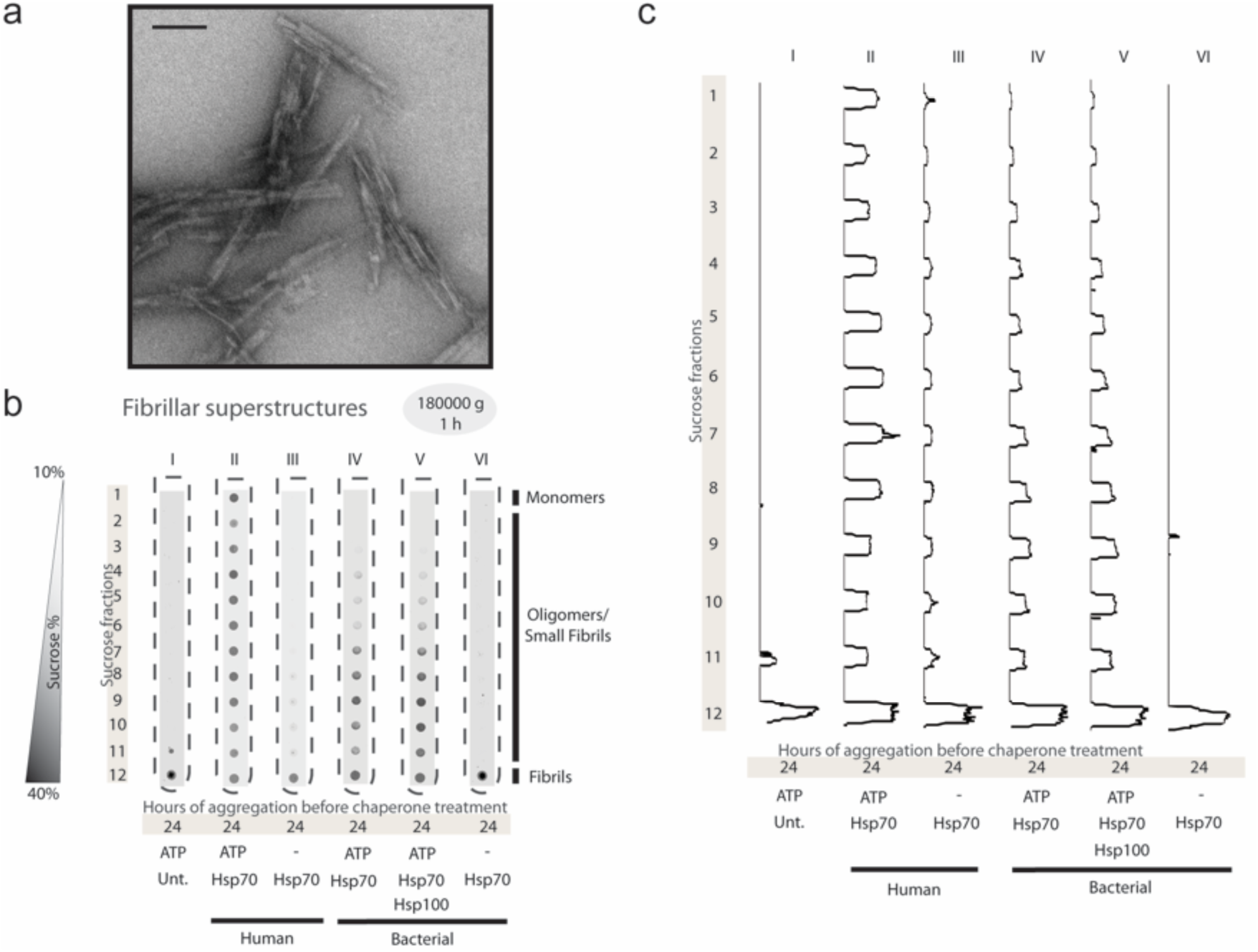
Disaggregation of Tau fibrillar superstructures by molecular chaperones. **a**. TEM of Tau fibrillar superstructures. Scale bar 100 nm. **b**. Density gradients of preformed, heparin-induced Tau-RD* fibrillar superstructures, untreated (Unt.) or treated for 24 h (human system) or 2 h (bacterial system) with chaperones as indicated. Centrifugation was performed at 185,000 g for 1 h. **c**. Intensity profiling of (**b**), normalized to a 0 to 1 scale.

In disease, Tau is part of larger tangles. We wondered whether association of Tau into such larger superstructures might affect the ability of Hsc70 to disassemble fibrils. To this end we generated superstructures by growing fibrils at higher ionic strength. TEM images confirmed that under these conditions Tau fibrils form larger bundles (**Fig. 2a** and **Fig. S2**). We monitored Hsc70-dependent disassembly of these superstructures by density gradient (**Fig. 2b and c**). Compared to the previous experiments on single fibrils we used smaller g-force and centrifuged the gradients for shorter time to optimise resolution for oligomers. Interestingly, the human Hsc70 system also disassembled the Tau superstructures into a mixture of monomers and oligomers in an ATP-dependent manner (**Fig. 2b**, tubes II and III). Again, after Hsc70 action we did not observe defined particles anymore by TEM and residual protein material became positively stained (**Fig. S2**). Thus, human Hsc70 can disassemble Tau fibrils, independent of their superstructure.

The Hsp70 system is conserved in evolution. The bacterial DnaK, together with its co-chaperones, the J-protein DnaJ and the nucleotide exchange factor GrpE, can disaggregate small soluble aggregates. In co-operation with the Hsp100 disaggregase ClpB, lacking in the metazoan cytosol, DnaK also disaggregates large, amorphous aggregates ^8,25^. Notably, Hsp100 disaggregases are lacking in the metazoan cytosol. However, DnaK and ClpB alone do not disaggregate Tau-RD fibrils ^9^. We confirmed that the bacterial system did not have any impact on single Tau fibrils (**Fig. 1b**, tubes V and VI; **Fig. 1c; Fig. S1a**). Remarkably, however, when applying the bacterial Hsp70 system to fibril superstructures we noted that the superstructures were dissolved (**Fig. 2b**, tubes IV and VI; **Fig. S2**). Thus, the *E. coli* Hsp70 system can disassemble the fibril superstructures, but it cannot disassemble the fibrils themselves. We conclude that nicking of Tau fibrils is an activity only acquired for Hsp70 systems later in evolution. Notably, Hsp100 did not even contribute further to disaggregation of fibril superstructures (**Fig. 2b**, tubes IV and V; **Fig. 2c**). We conclude that the bacterial Hsp70 and Hsp100 disaggregases lack the ability to disassemble single fibrils, despite their ability to effectively dissolve amorphous aggregates.

## DISCUSSION

Our findings suggest that the disaggregation activity of Hsp70 consists of two capacities – one linked to controlling fibril superstructure, which is an evolutionary conserved Hsp70 activity, the other linked to disaggregation of the fibril itself, which is reserved to the human homolog (**Fig. 3**). Thus, the capacity to disaggregate Tau fibrils is a unique property of the human system. The bacterial Hsp70/Hsp100 systems are unable to dissolve fibrils, although they are highly active on amorphous aggregates. In yeast, the Hsp100 but not the Hsp70 shows disaggregating activity for fibrils, which may be correlated with the presence of prions in yeast ^9, 26, 27^. This suggests that defibrillation is a meaningful Hsp70 activity acquired in evolution together with the increasing appearance of pro-fibrillic proteins. Interestingly, metazoa lack cytoplasmatic Hsp100 disaggregases. Thus, the ability to disaggregate amyloid structures by the Hsp70 machinery may have co-evolved in metazoa at the expense of the Hsp100 disaggregases.

**Figure 3.**
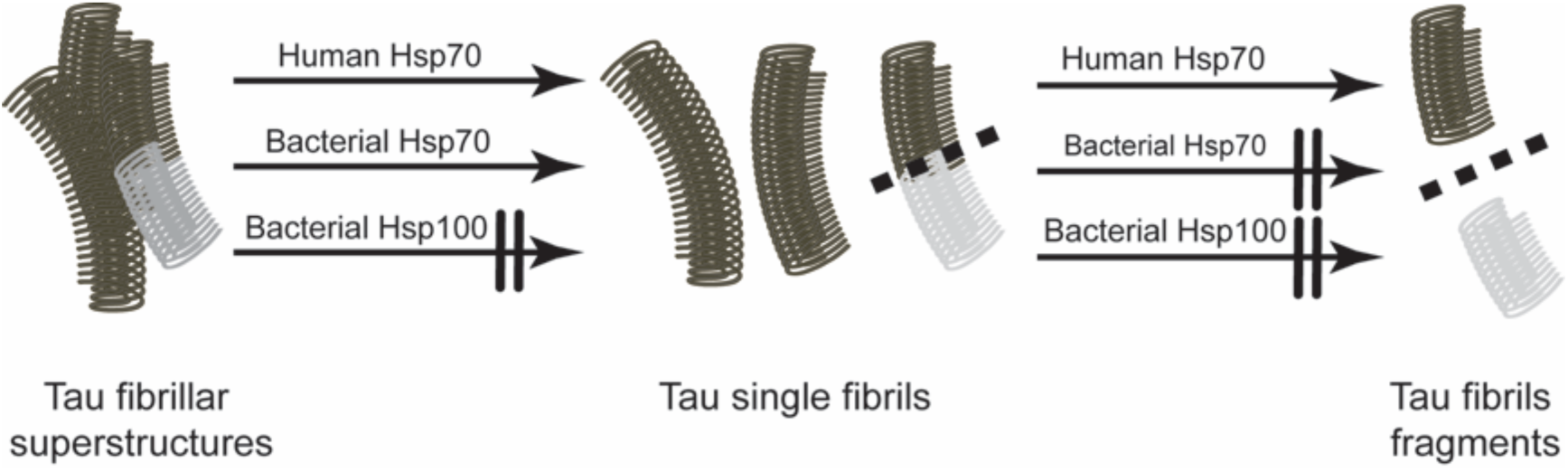
Unique adaptation of human Hsp70 to Tau fibrils. Only the human Hsp70 system can effectively disaggregate Tau tangles and single fibrils, suggesting fine-tuning of the disaggregation machineries during the co-evolution of chaperones and amyloid structures.

The ability of the Hsp70 system to dissolve neurotoxic fibrils of both Tau and α-synuclein endows human neurons with chaperone machinery capable of processing various amyloids. This is potentially relevant for other fibril diseases such as ALS or corea Huntington. It may be this fibril dissolving power that can also control liquid-liquid phase separation. Inhibiting Hsp70 increases the half-life of stress granules, pointing indirectly a role of this chaperone in the disassembly of stress granules ^28^.

Hsp70 is a central node of the proteostasis network, which gradually collapses during aging ^29, 30^. While it is established that the chaperone network controls protein homeostasis of monomeric Tau, our data show that Hsc70 has the ability to approach Tau after initiation of aggregation ^16-19^. This activity may be a double-edged sword. It may contribute to successful prevention of fibril formation for the first decades of our life, but it remains to be seen whether Hsc70 could contribute to generate potentially toxic oligomers from apparently inert fibrils. A natural disaggregase in neurons may be an interesting target for novel drug strategies in Alzheimer. Thus, It is an exciting challenge to control function of the Hsp70 machinery in the later stages of life.

## METHODS

### Protein purification

#### Purification of Tau-RD*

We overproduced N-terminally FLAG-tagged (DYKDDDDK) Human Tau-RD (Q244-E372, also referred to as K18, with pro-aggregation mutation ΔK280) recombinantly in *E. coli* BL21 Rosetta 2 (Novagen), with additional removable N-terminal His_6_-Smt-tag (MGHHHHHHGSDSEVNQEAKPEVKPEVKPETHINLKV SDGSSEIFFKIKKTTPLRRLMEAFAKRQGKEMDSLRFLYDGIRIQADQTP EDLDMEDNDIIEAHREQIGG). Cells were harvested, flash-frozen in liquid nitrogen and stored at -80°C until further usage. Pellets were thawed in a water bath at 37°C and resuspended in 50 mM HEPES-KOH pH 8.5 (Sigma-Aldrich), 50 mM KCl (Sigma-Aldrich), ½ tablet/50 ml EDTA-free protease inhibitor (Roche), 5 mM β-mercaptoethanol (Sigma-Aldrich). Cells were disrupted by an EmulsiFlex-C5 cell disruptor (Avestin). Lysate was cleared by centrifugation, filtered with a 0.22 μm polypropylene filter (VWR) and supernatant was purified using an ÄKTA Purifier chromatography system (GE Healthcare). First, protein was loaded onto a POROS 20MC (Thermo Fischer Scientific) affinity purification column in 50 mM HEPES-KOH pH 8.5, 50 mM KCl, eluted with a 0-100% linear gradient (5 Column Volumes, CV) of 1 M imidazole. Fractions of interest were collected and concentrated in a buffer concentrator (Vivaspin, cut-off 10 kDa) to final volume of 3 ml. The concentrated sample was desalted with a PD-10 desalting column (GH Healthcare) in 50 mM HEPES-KOH pH 8.5, ½ tablet/50 ml Complete protease inhibitor (Roche)) and 5 mM β-mercaptoethanol. The His_6_-Smt tag was removed by Ulp1 treatment, shaking at 4°C over night. Next day, protein was loaded onto a POROS 20HS (Thermo Fischer Scientific) cation exchange column equilibrated with 50 mM HEPES-KOH pH 8.5. Protein was eluted with a 0-100% linear gradient (15 CV) of 2 M KCl (Carl Roth). Fractions of interest were collected and loaded onto a HiLoad 26/60 Superdex 200 pg (GE Healthcare Life Sciences) size exclusion column equilibrated with 25 mM HEPES-KOH pH 7.5, 75 mM KCl, 75 mM NaCl and 10 mM DTT. Fractions of interest were further concentrated to the desired final concentration using a concentrator (Vivaspin, cut-off 5 kDa). Protein concentration was measured using an ND-1000 UV/Vis spectrophotometer (Nanodrop Technologies) and purity was assessed with SDS-PAGE. Protein was aliquoted and stored at -80°C.

#### Protein purification of DnaK

E. *coli* DnaK was purified by adapting a published procedure ^31, 32^. We overproduced N-terminally His_6_-Smt3-tagged bacterial DnaK recombinantly in ΔdnaK52 cells (BB1994). Cell pellets were resuspended in lysis buffer (20 mM Tris-HCl pH 7.9, 100 mM KCl, 1 mM PMSF) and disrupted by an EmulsiFlex-C5 cell disruptor. Lysate was cleared by centrifugation, filtered with a 0.22 μm polypropylene filter and loaded onto a column with Ni-IDA resin (Macherey-Nagel). Column was washed with 20 CV of lysis buffer, 10 CV of ATP buffer (20 mM Tris-HCl pH 7.9, 100 mM KCl, 5mM MgCl2, 5 mM ATP), and 2 CV of lysis buffer. Protein was eluted with 20 mM Tris-HCl pH 7.9, 100 mM KCl, 250 mM imidazole. The His_6_-Smt3 tag was removed by Ulp1 treatment. After cleavage and dialysis against lysis buffer, the protein mixture was loaded onto a Ni-IDA column and the flow-through fraction containing tag-free DnaK was collected. DnaK was loaded onto a Resource Q (GE Healthcare) anion exchange column equilibrated with 40 mM HEPES-KOH pH 7.6, 100 mM KCl, 5 mM MgCl2. DnaK was eluted with a 0.1-1 M linear gradient (10 CV) of KCl. The protein concentration was measured using an ND-1000 UV/Vis spectrophotometer and purity was assessed with SDS-PAGE. Protein was aliquoted and stored at -80°C.

#### Protein purification of DnaJ

*E. coli* DnaJ was purified by adapting a published procedure ^32, 33^. We overproduced DnaJ in *E. coli* strain W3110. Cell pellets were resuspended in lysis buffer (50 mM Tris-HCl pH 8, 10 mM DTT, 0.6% (w/v) Brij 58, 1 mM PMSF, 0.8 g/l Lysozyme) and disrupted by an EmulsiFlex-C5 cell disruptor. Lysate was cleared by centrifugation and filtered with a 0.22 μm polypropylene filter. One volume of buffer A (50 mM Sodium Phosphate pH 7, 5 mM DTT, 1 mM EDTA, 0.1% (w/v) Brij 58) was added to the supernatant and protein was precipitated by addition of (NH4)_2_SO_4_ to a final concentration of 65% (w/v). Protein was pelleted by centrifugation (15,000 g, 30 min), pellet was dissolved in 220 mL buffer B (50 mM Sodium Phosphate pH 7, 5 mM DTT, 1 mM EDTA, 0.1% (w/v) Brij 58, 2 M Urea) and dialysed against buffer B. DnaJ was then loaded onto a POROS 20HS cation exchange column equilibrated with buffer B, washed with buffer B and eluted with a 0-0.66 linear gradient (15 CV) of KCl. Fractions of interest were pooled and dialysed against buffer C (50 mM Tris-HCl pH 7.5, 2 M urea, 0.1% (w/v) Brij 58, 5 mM DTT, 50 mM KCl), then the sample was loaded onto a hydroxyapatite column equilibrated with buffer C. The column was first washed with 1 CV of buffer C supplemented with 1 M KCl, and then with 2 CV of buffer C. DnaJ was eluted with a 0%–50% linear gradient (1 CV) of buffer D (50 mM Tris-HCl pH 7.5, 2 M urea, 0.1% (w/v) Brij 58, 5 mM DTT, 50 mM KCl, 600 mM KH_2_PO_4_) and 2 CV of 50% buffer D. Fractions of interest were pooled and dialysed against buffer E (50 mM Tris-HCl pH 7.7, 100 mM KCl). Protein concentration was measured using an ND-1000 UV/Vis spectrophotometer and purity was assessed with SDS-PAGE. Protein was aliquoted and stored at -80°C.

#### Protein purification of GrpE

*E. coli* GrpE was purified by adapting a published procedure ^32, 34^. We overproduced GrpE in ΔdnaK52 cells (BB1994). Cell pellets were resuspended in lysis buffer (50 mM Tris-HCl pH 7.5, 100 mM KCl, 3 mM EDTA, 1 mM PMSF) and disrupted by an EmulsiFlex-C5 cell disruptor. Lysate was cleared by centrifugation and filtered with a 0.22 μm polypropylene filter. Protein was precipitated by addition of (NH4)_2_SO_4_ to a final concentration of 0.35 g/ml. Protein was pelleted by centrifugation (15,000 g, 30 min), pellet was dissolved in 200 mL buffer A (50 mM Tris-HCl pH 7.5, 100 mM KCl, 1 mM DTT, 1 mM EDTA, 10% glycerol) and dialysed twice against the same buffer for 4 h. Protein was loaded onto a HiTrap Q XL anion exchange column equilibrated with buffer A and then eluted using a linear gradient of buffer B (50 mM Tris-HCl pH 7.5, 1 M KCl, 1 mM DTT, 1 mM EDTA, 10% glycerol). Fractions of interest were dialysed against buffer C (10 mM Potassium Phosphate pH 6.8, 1 mM DTT, 10% glycerol). GrpE was loaded onto a Superdex 200 pg size exclusion column equilibrated with buffer A and concentrated using a HiTrap Q XL anion exchange column with a steep gradient. Protein concentration was measured using an ND-1000 UV/Vis spectrophotometer and purity was assessed with SDS-PAGE. Protein was aliquoted and stored at -80°C.

#### Protein purification of Hsc70

Human Hsc70 (gene name: HSPA8) was purified by adapting a published procedure ^35^. We overproduced N-terminally His_6_-Smt3-tagged Hsc70 recombinantly in *E. coli* BL21 Rosetta 2. Cell pellets were resuspended in lysis buffer (20 mM Tris-HCl pH 7.9, 100 mM KCl, ½ tablet/50 ml EDTA-free protease inhibitor) and disrupted by an EmulsiFlex-C5 cell disruptor. Lysate was cleared by centrifugation, filtered with a 0.22 μm polypropylene filter and supernatant was loaded onto a column with Ni-IDA resin. Column was washed with 20 CV of lysis buffer (Complete Protease Inhibitor instead of the EDTA-free one), 20 CV of high salt buffer (20 mM Tris-HCl pH 7.9, 1 M KCl) and again with 2 CV of lysis buffer. The column was then slowly washed with 10 CV of ATP-buffer (40 mM Tris-HCl pH 7.9, 100 mM KCl, 5 mM MgCl_2_, 5 mM ATP) to elute bound substrates. Protein was eluted with two times 1 CV of elution buffer (40 mM Tris-HCl pH 7.9, 100 mM KCl, 250 mM imidazole). Ulp1 was added to the elution and sample was dialyzed overnight against dialysis buffer (40 mM HEPES-KOH pH 7.6, 10 mM KCl, 5 mM MgCl_2_). Protein was then loaded on Ni-IDA material, flow-through containing Hsc70 was collected and loaded onto a POROS 20HQ (Thermo Fischer Scientific) cation exchange column. Hsp70 was eluted with a linear gradient of elution buffer (40 mM HEPES-KOH pH 7.6, 1 M KCl, 5 mM MgCl2, 10 mM μ-mercaptoethanol, 5% glycerol) and dialysed against 40 mM HEPES-KOH pH 7.6, 50 mM KCl, 5 mM MgCl_2_, 10 mM β-mercaptoethanol, 10% glycerol. Protein concentration was measured using an ND-1000 UV/Vis spectrophotometer and purity was assessed with SDS-PAGE. Protein was aliquoted and stored at -80°C.

#### Protein purification of Hdj1

Human Hdj1 (gene name: DNAJB1) was purified by adapting a published procedure ^32, 36^. We overproduced N-terminally His_6_-Smt3-tagged human Hdj1 recombinantly in in *E. coli* BL21 Rosetta 2 strain. Cell pellets were resuspended in lysis buffer (20 mM Tris-HCl 20 pH 7.9, 100 mM KCl, ½ tablet/50 ml EDTA-free protease inhibitor) and disrupted by an EmulsiFlex-C5 cell disruptor. Lysate was cleared by centrifugation, filtered with a 0.22 μm polypropylene filter, loaded onto a POROS 20MC affinity purification column equilibrated with lysis buffer, then eluted with a 0-100% linear gradient (5 CV) of 1 M imidazole. The His_6_-Smt tag was removed by Ulp1 treatment. After cleavage and dialysis against 25 mM HEPES-KOH, 350 mM KCl, 5 mM MgCl_2_, 5 mM β-mercaptoethanol, the protein mixture was loaded onto a POROS 20HS cation exchange column equilibrated with 50 mM HEPES-KOH pH 8.5. Protein was eluted with a 0-100% linear gradient (15 CV) of 2 M KCl. Fractions of interest were buffer exchanged against 25 mM HEPES-KOH pH 7.5, 75 mM KCl, 75 mM NaCl and 10 mM DTT, then further concentrated using a concentrator. Protein concentration was measured using an ND-1000 UV/Vis spectrophotometer and purity was assessed with SDS-PAGE. Protein was aliquoted and stored at -80°C.

#### Protein purification of Apg2

Human Apg2 was purified by adapting a published procedure ^32, 35^. We overproduced N-terminally His_6_-Smt3-tagged human Apg2 recombinantly in in *E. coli* BL21 Rosetta 2 strain. Cell pellets were resuspended in lysis buffer (40 mM Tris-HCl pH 7.9, 100 mM KCl, 5 mM ATP, ½ tablet/50 ml EDTA-free protease inhibitor) and disrupted by an EmulsiFlex-C5 cell disruptor. Lysate was cleared by centrifugation, filtered with a 0.22 μm polypropylene filter and loaded onto a column with Ni-IDA resin. Column was washed with 20 CV of wash buffer (40 mM Tris-HCl pH 7.9, 100 mM KCl, 5 mM ATP) followed by elution with elution buffer (40 mM Tris-HCl pH 7.9, 100 mM KCl, 300 mM imidazole). Protein was then loaded on a HiPrep 26/10 Desalting Column equilibrated with 20 mM Tris-HCl pH 7.9, 100 mM KCl, 10% glycerol. The His_6_-Smt tag was removed by Ulp1 shaking at 4°C over night, in the presence of 5 mM ATP. The day after, protein was loaded onto a HiLoad 16/600 Superdex 200pg size exclusion column equilibrated with 40 mM HEPES-KOH pH 7.6, 10 mM KCl, 5 mM MgCl2, 10% glycerol. Fractions of interest were loaded onto a Resource Q anion exchange column equilibrated with 40 mM HEPES-KOH pH 7.6, 10 mM KCl, 5 mM MgCl2, 10% glycerol. Protein was eluted with a 0-100% linear gradient (10 CV) of 1 M KCl. Protein concentration was measured using an ND-1000 UV/Vis spectrophotometer and purity was assessed with SDS-PAGE. Protein was aliquoted and stored at -80°C.

#### Protein purification of ClpB

*E. coli* ClpB was purified by adapting a published procedure ^8^. We overproduced C-terminally His_6_-tagged bacterial ClpB recombinantly in *E. coli* XI1-blue cells. Cell pellets were resuspended in lysis buffer (20 mM Tris-HCl pH 7.9, 100 mM KCl, ½ tablet/50 ml EDTA-free protease inhibitor) and disrupted by an EmulsiFlex-C5 cell disruptor. Lysate was cleared by centrifugation, filtered with a 0.22 μm polypropylene filter, loaded onto an POROS 20MC affinity purification column equilibrated with lysis buffer, then eluted with a 0-100% linear gradient (5 CV) of 1 M imidazole. Fractions of interest were collected and loaded onto a HiLoad 26/60 Superdex 200 pg size exclusion column equilibrated with buffer 25 mM HEPES-KOH pH 7.5, 75 mM KCl, 75 mM NaCl and 10 mM DTT. Fractions of interest were further concentrated to the desired final concentration using a concentrator. Protein concentration was measured using an ND-1000 UV/Vis spectrophotometer and purity was assessed with SDS-PAGE. Protein was aliquoted and stored at -80°C.

### Protein aggregates

Monomeric Tau-RD* (10 *μ*M) was aggregated in 25 mM HEPES-KOH pH 7.5 with Complete protease inhibitor (1/2 tablet/50 ml), DTT 10 mM (Thermo-Scientific) and 2.5 *μ*M of Heparin (Santa Cruz Biotech). Fibrils were aliquoted and flash frozen after 24 h of aggregation at 37°C, shaking. Single fibrils were obtained by in aggregation buffer containing 75 mM KCl / 75 mM NaCl, whereas fibrillar superstructures were obtained in aggregation buffer containing 500 mM KCl / 75 mM NaCl.

### Disaggregation experiments

Chaperones mix (10 *μ*M Hsc70, 5 *μ*M Hdj1, 2,3 *μ*M Apg2 for human system; 10 *μ*M DnaK, 2 *μ*M DnaJ and 5 *μ*M GrpE for bacterial system) and Tau fibrils (Tau monomeric concentration: 2.5 *μ*M) were incubated at 37°C, shaking at 180 rpm. Incubation time for each experiment is specified in the text. After incubation, samples were loaded completely on top of density gradients composed of a 10 to 40% sucrose gradient. Gradients were run on a Sorvall WX Ultra Series Centrifuge WX80 (Thermo Scientific), time and centrifugal forces depending on the experiment. Fractions were collected manually, each fraction 1/12^th^ of the total volume.

### Transmission Electron Microscopy

Specimens were prepared for Transmission Electron Microscopy using a negative staining procedure. Briefly, a 5 *μ*l drop of sample solution was adsorbed to a glow-discharged (twice for 20 s, on a Kensington carbon coater) pioloform-coated copper grid, washed five times on drops of deionised water, and stained with two drops of freshly prepared 2.0% Uranyl Acetate, for 1 and 5 minutes, respectively, and subsequently air dried. Samples were imaged at room temperature using a Tecnai T20 LaB_6_ electron microscope operated at an acceleration voltage of 200 kV and equipped with a slow-scan Gatan 4K x 4K CCD camera. Images were acquired at a defocus value of 1.5 *μ*m. Magnification values and pixel sizes on specimen level are summarised in Supplementary Table S1.

### Production of density gradients

Density gradients were formed by dissolving 10% and 40% sucrose (Sigma-Aldrich), in 25 mM HEPES pH 7.5, 75 mM KCl, 75 mM NaCl. Gradients were set up in polyallomer centrifuge tubes (Beckmann) by filling them to half volume with 40% sucrose and topping them up with an equal amount of 10% sucrose. Gradients were formed by tilting the tubes horizontally for 3 h at room temperate and then tilting them back to vertical position. Tubes were stored overnight at 4°C.

### Dot blot analysis of density gradients

Dot blot was performed using a dot blot apparatus (BioRad) and nitrocellulose membrane 0.1 *μ*M (Sigma-Aldrich) washed with PBS. Each well was filled with 150 *μ*L of sample, incubated on the membrane 10 minutes at room temperature and then pulled through by applying vacuum. Nitrocellulose membranes were blocked with PBS blocking buffer (LI-COR) at room temperature for 1 hour and then incubated with primary antibody monoclonal anti-FLAG M2 (F3165, Sigma Aldrich, working _dilution_ 1:1000) at room temperature for 1 hour. After three washes with PBS, secondary antibody Donkey anti Mouse IgG IR Dye 800 conjugated (610-732-002, Rockland, 1:5000) was added at room temperature for 45 minutes. After additional 2 washes with PBS-T and one final wash with PBS, detection was performed using Odyssey CLx (LI-COR). Quantification was performed via Image Studio Lite (LI-COR).

### Dot blot profiling

For each tube, a rectangle encompassing all 12 fractions was drawn. The rectangle height corresponded to half diameter of each dots. Profiles were obtained via ImageJ software. Raw data was normalized to 0-1 range and plotted. The presence of visible aggregates in higher fractions precluded linear interpretation, as larger particles affect both accessibility for the antibody and the fluorescence signal itself.

## ACKNOWLEDGMENTS

We are grateful to Ineke Braakman for continuous support. We thank Axel Mogk for kindly providing us with a bacterial stab of *E. coli* XL1 pClpB. We thank Madelon Maurice for collaboration in the Initial Training Network “WntsApp” (No. 608180)], supported by Marie-Curie Actions of the 7th Framework programme of the EU. SGDR was further supported by the Internationale Stichting Alzheimer Onderzoek (ISAO; project “Chaperoning Tau Aggregation”; No. 14542) and a ZonMW TOP grant (“Chaperoning Axonal Transport in neurodegenerative disease”; No. 91215084).

## AUTHOR CONTRIBUTIONS

S.G.D.R. and L.F. conceived the study; S.G.D.R., L.F. and F.G.F. planned experiments; L.F., W.J.C.G., M.v.W., R.K., K.K. and L.v.B. did experiments; L.F., W.J.C.G., M.v.W., R.K., K.K. and L.v.B. analysed data; S.G.D.R. and L.F. wrote the manuscript, with contributions of all authors.

## MATERIALS AND CORRESPONDENCE

Correspondence and material requests should be addressed to S.G.D.R.

## SUPPLEMENTARY MATERIAL

**SUPPLEMENTARY TABLE S1.** Magnification values and pixel sizes on specimen level for all TEM images.

| Figure | Magnification | nm / pixel |
| --- | --- | --- |
| Fig. 1A | 62,000 | 0.178 |
| Fig. 2A | 62,000 | 0.178 |
| Fig. S1a, left panel up | 11,500 | 0.954 |
| Fig. S1a, mid panel up | 14,500 | 0.746 |
| Fig. S1a, right panel up | 6,500 | 1.681 |
| Fig. S1a, left panel down | 62,000 | 0.178 |
| Fig. S1a, mid panel down | 29,000 | 0.37 |
| Fig. S1a, right panel down | 62,000 | 0.178 |
| Fig. S1b | 62,000 | 0.18 |
| Fig. S2, left panel up | 29,000 | 0.37 |
| FigS2, mid panel up | 19,000 | 0.561 |
| FigS2, right panel up | 14,500 | 0.746 |
| FigS2, left panel down | 62,000 | 0.178 |
| FigS2, mid panel down | 19,000 | 0.561 |
| FigS2, right panel down | 62,000 | 0.178 |

**Supplementary Figure 1.**
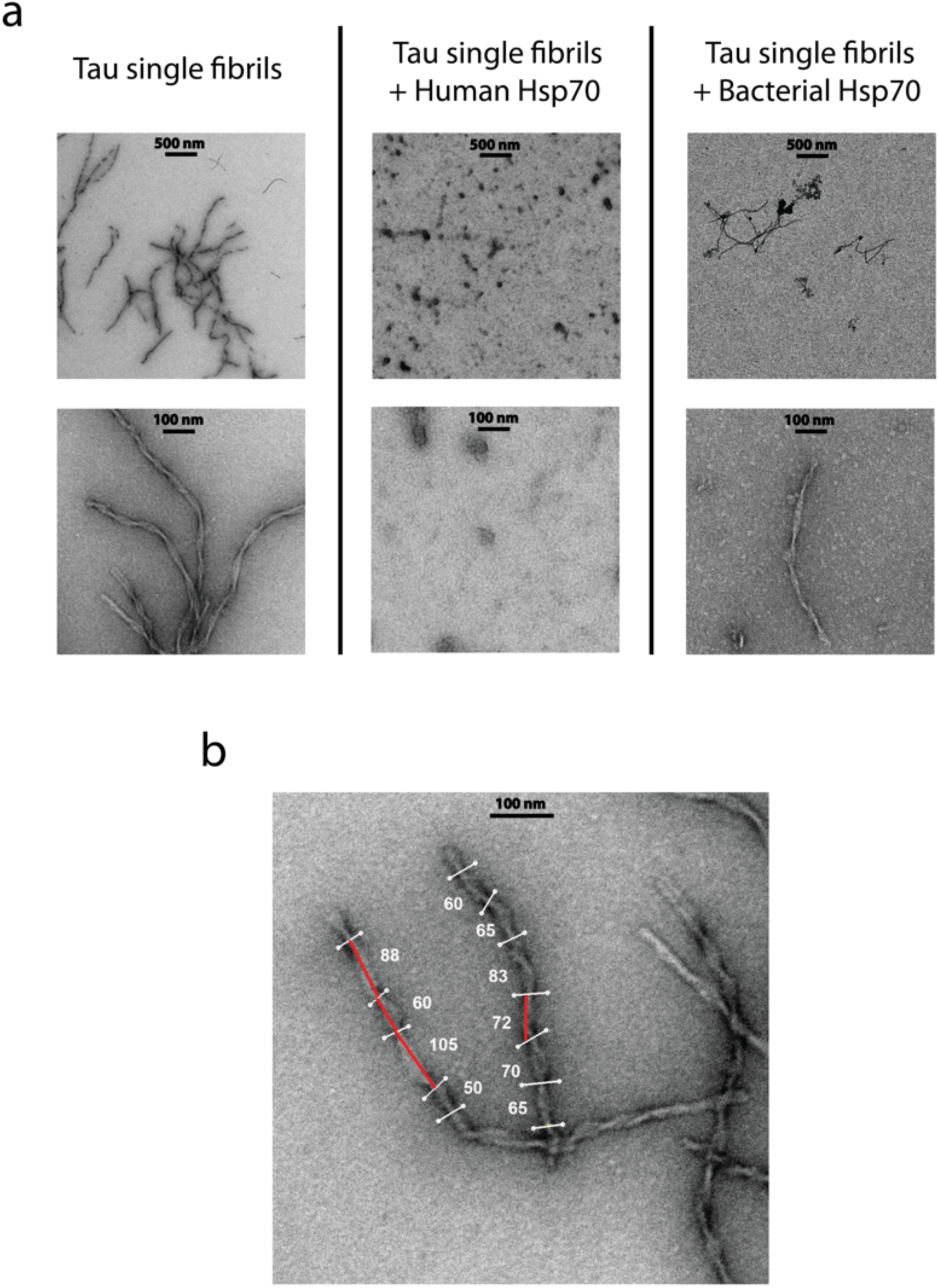
Additional TEM images of human chaperones disaggregating Tau single fibrils, bacterial chaperones incapable of disaggregation. **a**. TEM of Tau single fibrils, untreated or treated for 24 h with chaperones as indicated. Residual protein material after treatment with human chaperones became positively stained. Scale bars vary as indicated. **b**. Estimation of crossover distances for typical Tau fibrils (numbers indicate crossover distances in nm).

**Supplementary Figure S2.**
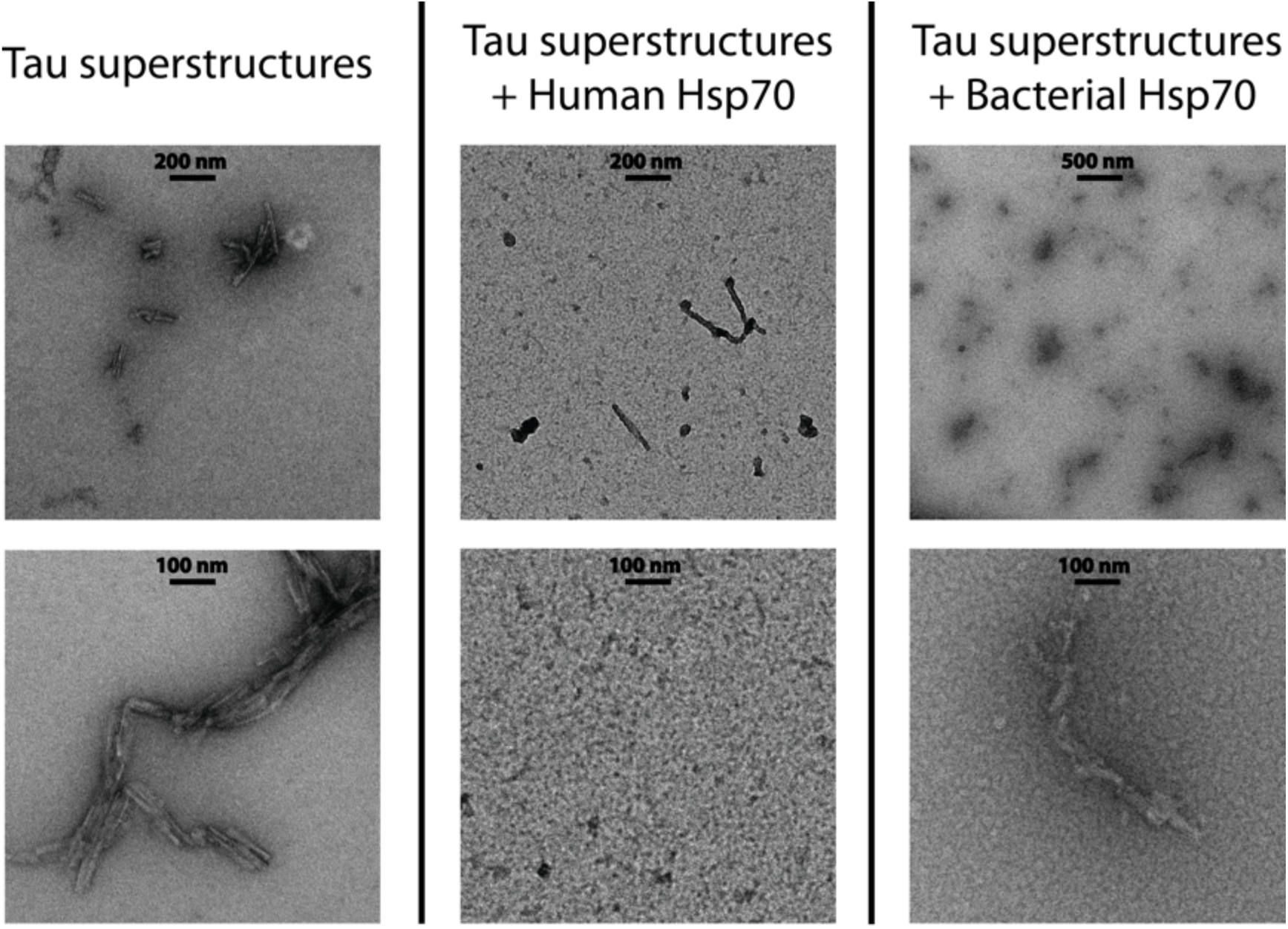
Additional TEM images of human and bacterial chaperones disaggregating Tau fibrillar superstructures. TEM of Tau single fibrils, untreated or treated for 24 h (human system) or 2 h (bacterial system) with chaperones as indicated. Residual protein material after treatment with human chaperones became positively stained. Scale bars vary as indicated.

## References

1. Wilcock, G. K. & Esiri, M. M. Plaques, tangles and dementia. A quantitative study. J. Neurol. Sci. 56, 343–356 (1982).

2. Holtzman, D. M. et al. Tau: From research to clinical development. Alzheimers Dement. 12, 1033–1039 (2016).

3. Wang, Y. & Mandelkow, E. Tau in physiology and pathology. Nat. Rev. Neurosci. 17, 5–21 (2016).

4. Goedert, M., Eisenberg, D. S. & Crowther, R. A. Propagation of Tau Aggregates and Neurodegeneration. Annu. Rev. Neurosci. 40, 189–210 (2017).

5. Hartl, F. U. Protein Misfolding Diseases. Annu. Rev. Biochem. (2017).

6. Stoecklin, G. & Bukau, B. Telling right from wrong in life - cellular quality control. Nat. Rev. Mol. Cell Biol. 14, 613–615 (2013).

7. Mogk, A., Kummer, E. & Bukau, B. Cooperation of Hsp70 and Hsp100 chaperone machines in protein disaggregation. Front. Mol. Biosci. 2, 22 (2015).

8. Mogk, A. et al. Identification of thermolabile Escherichia coli proteins: prevention and reversion of aggregation by DnaK and ClpB. Embo J 18, 6934–49 (1999).

9. DeSantis, M. E. et al. Operational plasticity enables hsp104 to disaggregate diverse amyloid and nonamyloid clients. Cell 151, 778–793 (2012).

10. Mukrasch, M. D. et al. Structural Polymorphism of 441-Residue Tau at Single Residue Resolution. Plos Biology 7, 399–414 (2009).

11. Gustke, N., Trinczek, B., Biernat, J., Mandelkow, E. M. & Mandelkow, E. Domains of Tau-Protein and Interactions with Microtubules. Biochemistry (N. Y.) 33, 9511–9522 (1994).

12. Sato, C. et al. Tau Kinetics in Neurons and the Human Central Nervous System. Neuron 97, 1284–1298.e7 (2018).

13. Fitzpatrick, A. W. P. et al. Cryo-EM structures of tau filaments from Alzheimer's disease. Nature 547, 185–190 (2017).

14. Falcon, B. et al. Structures of filaments from Pick's disease reveal a novel tau protein fold. Nature 561, 137–140 (2018).

15. Mocanu, M. M. et al. The potential for beta-structure in the repeat domain of Tau protein determines aggregation, synaptic decay, neuronal loss, and coassembly with endogenous Tau in inducible mouse models of tauopathy. Journal of Neuroscience 28, 737–748 (2008).

16. Fontaine, S. N. et al. Cellular factors modulating the mechanism of tau protein aggregation. Cell Mol. Life Sci. 72, 1863–1879 (2015).

17. Karagöz, G. E. et al. Hsp90-Tau complex reveals molecular basis for specificity in chaperone action. Cell 156, 963–974 (2014).

18. Fontaine, S. N. et al. The active Hsc70/tau complex can be exploited to enhance tau turnover without damaging microtubule dynamics. Hum. Mol. Genet. 24, 3971–3981 (2015).

19. Dickey, C. A. et al. The high-affinity HSP90-CHIP complex recognizes and selectively degrades phosphorylated tau client proteins. J. Clin. Invest. 117, 648–658 (2007).

20. Nillegoda, N. B. et al. Crucial HSP70 co-chaperone complex unlocks metazoan protein disaggregation. Nature 524, 247–251 (2015).

21. Gao, X. et al. Human Hsp70 Disaggregase Reverses Parkinson's-Linked alpha-Synuclein Amyloid Fibrils. Mol. Cell (2015).

22. Sarkar, M., Kuret, J. & Lee, G. Two motifs within the tau microtubule-binding domain mediate its association with the hsc70 molecular chaperone. J. Neurosci. Res. 86, 2763–2773 (2008).

23. Jinwal, U. K. et al. Imbalance of Hsp70 family variants fosters tau accumulation. FASEB J. (2012).

24. Goedert, M. et al. Assembly of microtubule-associated protein tau into Alzheimer-like filaments induced by sulphated glycosaminoglycans. Nature 383, 550–553 (1996).

25. Goloubinoff, P., Mogk, A., Zvi, A. P., Tomoyasu, T. & Bukau, B. Sequential mechanism of solubilization and refolding of stable protein aggregates by a bichaperone network. Proc. Natl. Acad. Sci. U. S. A. 96, 13732–13737 (1999).

26. Glover, J. R. & Lindquist, S. Hsp104, Hsp70, and Hsp40: a novel chaperone system that rescues previously aggregated proteins. Cell 94, 73–82 (1998).

27. Glover, J. R. et al. Self-seeded fibers formed by Sup35, the protein determinant of [PSI+], a heritable prion-like factor of S. cerevisiae. Cell 89, 811–819 (1997).

28. Alberti, S., Mateju, D., Mediani, L. & Carra, S. Granulostasis: Protein Quality Control of RNP Granules. Front. Mol. Neurosci. 10, 84 (2017).

29. Labbadia, J. & Morimoto, R. I. The biology of proteostasis in aging and disease. Annu. Rev. Biochem. 84, 435–464 (2015).

30. Klaips, C. L., Jayaraj, G. G. & Hartl, F. U. Pathways of cellular proteostasis in aging and disease. J. Cell Biol. 217, 51–63 (2018).

31. Kityk, R., Vogel, M., Schlecht, R., Bukau, B. & Mayer, M. P. Pathways of allosteric regulation in Hsp70 chaperones. Nat. Commun. 6, 8308 (2015).

32. Morán Luengo, T., Kityk, R., Mayer, M. P. & Rüdiger, S. G. D. Hsp90 Breaks the Deadlock of the Hsp70 Chaperone System. Mol. Cell 70, 545–552.e9 (2018).

33. Graf, C., Stankiewicz, M., Kramer, G. & Mayer, M. P. Spatially and kinetically resolved changes in the conformational dynamics of the Hsp90 chaperone machine. EMBO J. 28, 602–613 (2009).

34. Schönfeld, H. J., Schmidt, D., Schröder, H. & Bukau, B. The DnaK chaperone system of Escherichia coli: quaternary structures and interactions of the DnaK and GrpE components. J. Biol. Chem. 270, 2183–2189 (1995).

35. Andreasson, C., Fiaux, J., Rampelt, H., Mayer, M. P. & Bukau, B. Hsp110 Is a Nucleotide-activated Exchange Factor for Hsp70. Journal of Biological Chemistry} 283, 8877–8884 (2008).

36. Malakhov, M. P. et al. SUMO fusions and SUMO-specific protease for efficient expression and purification of proteins. J. Struct. Funct. Genomics 5, 75–86 (2004).

